# Axon Diameter Measurements using Diffusion MRI are Infeasible

**DOI:** 10.1101/2020.10.01.320507

**Authors:** Michael Paquette, Cornelius Eichner, Thomas R. Knösche, Alfred Anwander

## Abstract

The feasibility of non-invasive axonal diameter quantification with diffusion MRI is a strongly debated topic due to the neuroscientific potential of such information and its relevance for the axonal signal transmission speed. It has been shown that under ideal conditions, the minimal diameter producing detectable signal decay is bigger than most human axons in the brain, even using the strongest currently available MRI systems. We show that resolving the simplest situations including multiple diameters is unfeasible even with diameters much bigger than the diameter limit. Additionally, the recently proposed effective diameter resulting from fitting a single value over a distribution is almost exclusively influenced by the biggest axons. We show how impractical this metric is for comparing different distributions. Overall, axon diameters currently cannot be quantified by diffusion MRI in any relevant way.

## 1 Introduction

In-vivo estimation of axon diameters has been an important goal of many researchers since the inception of diffusion MRI. As the diameter of a myelinated axon is one of the main determiners of its signal transmission velocity [20, 44], the availability of this structural information would greatly facilitate the description and functional modeling of the brain communication pathways on an individual basis [45]. Detailed knowledge of tract-specific axonal diameters would provide insight into detailed and mechanistic relationships between brain structure and important aspects of brain function, including development and learning. The capacity of dMRI to noninvasively probe cellular and axonal boundaries at the micrometer level seemed a promising method to pursue this aim.

The impact of restricted incoherent motion of water molecules on diffusion-weighted NMR signals has already been described in the early days of MR spectroscopy [39, 51]. However, these models only describe the diffusion process happening in the perpendicular cross-section of the axon. Using them to approximate axonal diameters requires prior knowledge of the tissue orientations, an equal diameter of all axons in the probed volume, as well as the absence of extra-axonal signals. A common strategy to bring it to the in-vivo 3D acquisition setting has been to combine one or many cylindrical compartments, describing the intra-axonal diffusion, with additional compartments describing the extra-axonal Gaussian diffusion process [3, 6, 7, 16].

Despite the overestimation of axonal diameters arising from the use of multi-compartment models [23] compared to electron microscopy ground truth [1, 37], these models are still seen as promising by a part of the community. This dilemma can be attributed to the fact that the relative *trend* of fitted diameters was argued to be somewhat plausible across the different parts of the corpus callosum [3, 5, 23, 29, 30] and that multi-compartment models in dMRI are difficult to fit reliably as they are essentially weighted sums of exponential functions.

Recent work highlighted an unavoidable sensitivity issue for detecting axon diameters of realistic size in the human brain, even with the latest high-end MRI systems [21, 40]. It proposes an “axon diameter limit” (*d*_min_) which corresponds to the smallest diameter that can be differentiated from a stick of diameter zero for given sequence parameters under ideal conditions. This *d*_min_ is computed from the most generous setting and is therefore a lower bound on the unbiased smallest diameter detectable for data deviating from the idealized case of diffusion signal arising only from parallel cylinders of equal diameter. The diameter limit suggests that previous “trends” in the estimated diameters are not supported by the measured data. Indeed, not only is the expected signal decay for restricted diffusion in realistically sized human axons very small, but it is also insensitive to changes in the gradient spacing in time (Δ), which is typically the parameter varied when the “small-big-small diameter trend” of the corpus callosum is observed [5, 10, 29, 30]. The large signal decay observed could be caused by noise, errors in the compartment separation, or by other types of time-dependent diffusion such as diffusion signal from the extra-axonal compartment, which is sensitive to Δ.

In this work, we employ extensive simulations of restricted diffusion MRI measurements under optimal conditions to concretely showcase the limitations of axon diameter mapping. We first show the small magnitude of the signal decay that results from perpendicular diffusion in axons with realistic diameters. This result highlights the sensitivity required to distinguish this additional signal drop from the rest of the unrestricted diffusion within a voxel. Secondly, we show the error when fitting diameters to noisy data, even in ideal situations. This experiment numerically showcases the axon diameter limit [40] which highlights the poor scaling exponents between controllable parameters and achievable diameter limits. We then extend our simulations from the case of single diameter estimations to that of fitting distributions of axonal diameters. Naturally, the same sensitivity limitations are present and even amplified since each diameter only produces a fraction of the total signal decay. Additionally, fitting errors do not concentrate on the ground truth diameters. Finally, we highlight the difficulty of interpreting a single diameter value fitted over a distribution, the so-called effective diameter [14, 52]. The effective diameter formulation adequately captures the averaging mechanisms which mix the signal decay contribution of a distribution of axon sizes. However, the effective diameter still suffers from the same sensitivity and specificity issues as presented before. The limitations of MR axon diameter estimation discussed in this work affect every diffusion model fitting diameters, as they all rely on the capacity of dMRI to detect small signal decay from inside a cylinder compartment [3, 7, 16, 24, 25].

## 2 Methods

### 2.1 Relevant parameters

Throughout this work, we used numerical simulations to showcase the sensitivity of dMRI to axon diameters. It is therefore crucial to use realistic values for the various physical parameters. We describe each parameter, their realistic ranges, and our default choices. Particularly, we are concerned with the order of magnitude of the quantities and their scaling behavior (see eq. 1). For completion, we provide scripts to recompute any quantity, figure or experiment, for any choice of parameters (https://github.com/mpaquette/axDiamFig).

#### Axon (cylinder) diameter (*d*)

The smaller the diameter, the smaller is the maximal displacement of the water molecules, as we assume impermeable axonal walls. This restricted water diffusion perpendicular to the axon will induce a small signal change proportional to the mean squared displacement inside the circular cross-section. Prior results from histological assessments show that human axons in the white matter of the brain have diameters in the order of 1 *μ*m [1, 37]. Typical distributions of diameters tend to peak around 0.5-1.0 *μ*m with maximum axon diameters around 2.5-5 *μ*m (see fig 4). Informally, the minimum sensitivity required to properly qualify such distributions has to be smaller than the peak of the distribution.

#### Unrestricted diffusivity of the medium (*D*_0_)

The lower the diffusivity is, the more time it takes for the diffusion process to saturate inside of the restricted compartment. The value of in-vivo intra-axonal diffusivity is still an actively studied topic. Early models in the literature chose to fix the in-vivo intra-axonal water compartment diffusivity to 1.7 *μ*m^2^/ms [2, 55]. Many multi-compartment models have the ambiguity of allowing two distinct solutions, depending on if the intra-axonal diffusivity is higher than the parallel extra-axonal diffusivity [33]. Indeed, recent results seem to indicate that intra-axonal diffusivity is higher [34]. A recent approach, using a planar diffusion filter to eliminate extra-axonal signal, reports values around 2.25 *μ*m^2^/ms [19] which roughly matches with previous animal studies, when accounting for tissue temperature [12, 49]. In the case of post-mortem measurements, both the reduced tissue temperature and the fixation process reduce the tissue diffusivity [43]. Reported values for post-mortem diffusivities are around 1/3 - 1/4 of that of in-vivo [22]. In our simulations we assume the following diffusivities: *D*_0*,in-vivo*_ = 2 *μ*m^2^/ms (2 × 10^−9^ m^2^/s), and *D*_0*,post-mortem*_ = 0.66 *μ*m^2^/ms (0.66 × 10^−9^ m^2^/s).

#### Diffusion gradient magnitude (G)

The diffusion gradient hardware varies among the different types of MRI scanners. The strength of diffusion gradient directly affects the signal decay in diffusion, *i.e.* the same total displacement of water molecule produces a bigger signal decay with a stronger gradient. Typical clinical scanners tend to have weaker gradients (*G*_max_ = 40 mT/m), while gradient coils in preclinical small-bore scanners can produce magnetic field gradients as strong as 1500 mT/m. For human in-vivo measurements, the Siemens Connectom MRI scanner (Siemens Healthineers, Erlangen, Germany) is the system that produces by far the strongest diffusion gradients (*G*_max_ = 300 mT/m). In our simulations we use *G* = 300 mT/m, as one of the goals associated with the development of this specific MRI system was to enable in-vivo axon diameter estimation.

#### Diffusion gradient duration (δ)

In the relevant regimes for human axon diameter estimation, the duration of the diffusion gradient pulse *δ* is the parameter probing the time-dependent diffusivity of restricted diffusion. Indeed, with a typical short achievable gradient pulse duration of around 5 ms on a human MRI system, we are well into the regime where the gradient duration is comparable or above the saturation time of the restricted compartment. In this regime, longer gradient pulses increase sensitivity (see sec. A.1). We limit the simulations to *δ*_max_ = 40 ms as longer pulses are impractical, as they increase the echo times of the acquisition, resulting in additional signal losses.

#### Diffusion gradient separation (Δ)

In the relevant regimes for human axon diameter estimation, the diffusion process is already saturated during the gradient application, and varying the temporal separation of the diffusion gradient pulses Δ provides no extra sensitivity to restricted diffusion (see sec. A.1 and fig. 7). Therefore, to maximize the signal, we use Δ = *δ*. In practice, varying Δ *could* still be necessary for multi-compartment models where it is necessary to disentangle intra-and extra-axonal signal contributions.

#### Signal to noise ratio (SNR)

Ultimately, the SNR is the key parameter upon which “sensitivity” is defined. Throughout the simulations presented in this study, we corrupt signals with Gaussian noise (for simplicity and to produce a best-case scenario), *i.e. S*_*noisy*_ = *S*_*noiseless*_ + *ϵ* where 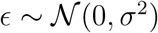. Since we only look at idealized diffusion effects, our signals have a value of 1 at the b0 (no diffusion gradient applied), and therefore the SNR is defined as SNR = *σ*^−1^. For comparison, the SNR of the b0 in the corpus callosum for a single in-vivo volume on the Connectom system with an echo time of 70 ms, repetition time 7500 ms, and resolution 1.8 mm isotropic is around 20. We showcase results for SNR = 30 and some results for SNR = 300, which correspond to 100 averages of a high-quality Connectom acquisition. Some diameter estimation approaches use aggregated fitting strategies such as region of interest (ROI) averaging or averaging along a tractography streamline path [4, 8, 9, 17] to increase the nominal SNR. These aggregated strategies make strong assumptions on tissue composition and orientation homogeneity in a region or along the entire pathway. It is unclear if the SNR gains of such strategies outweigh the bias due to tissue inhomogeneities in neighboring voxels as these methods still suffer from diameter overestimation [8].

#### Pulse Gradient Spin Echo Sequence (PGSE)

The PGSE sequence is maximally sensitive to perpendicular diffusion inside straight cylinders [40]. Therefore, we exclusively consider this sequence to showcase the sensitivity in the best-case scenario. When we consider axons misaligned with the gradient directions or undulating axons, other sequences such as oscillating gradient spin-echo (OGSE) can be more sensitive than PGSE [13, 36, 40, 41]. However, the key point is that any sequence in those scenarios is still less sensitive to the diameter than PGSE is in the non-misaligned and non-undulation case. Therefore, showing sensitivity issues with PGSE in that optimal case bounds all cases.

### 2.2 dMRI signal sensitivity to the diameter

Diffusion MRI contrast is related to the bulk displacement of the water molecules during the diffusion encoding, which causes the measured signal decay. Inside restricted compartments such as the cross-section of a cylinder, the maximal displacement is capped by the boundary, potentially producing much smaller signal decays than produced by free diffusion. These restricted diffusion processes can be classified into different time regimes. On short time scales, the bulk of water molecules has not yet interacted with the boundary and therefore behaves as in free diffusion. In the long time regime (wide pulse regime), most molecules have significantly interacted with the boundary and their position at any given time does not correlate with their initial position inside the cross-section of the axon; *i.e.* the signal has reached maximal decay from diffusion effects.

The general perpendicular signal decay formula for a cylinder using a Pulsed Gradient Spin Echo (PGSE) diffusion sequence [48] was first described by Neuman [39] and then extended by Van Gelderen [51] to account for cases where Δ ≠ *δ* (eq. 3). For the parameter ranges described in sec. 2.1, the Neuman long time limit (eq. 4) produces almost indistinguishable results. In this work, we use eq. 3 truncated to 50 terms to generate and fit signals arising from restricted diffusion.

For realistic acquisition and biological relevant parameter values (see sec 2.1), the diffusion process falls into the long time regime (wide pulse regime) and the expected signal decay is small compared to the noise amplitude at typical SNR. Using eq. 3, we simulated the expected MR signal decay for a multitude of combinations and we report the **decay percentage** values in Table 1. To cover a wide range of biological, experimental, and instrumental parameters, we simulated restricted diffusion MRI signals using (i) both in-vivo and post-mortem diffusivities, (ii) clinical gradient systems, and high-end Connectom gradients, and (iii) small to large human axon diameters.

**Table 1:** MR signal **decay** (in percent) for various diffusivities, acquisition parameters, and axon diameters. We note that if we have SNR = 30, a noise realization of one standard deviation has a magnitude 3.3% signal decay. This showcases the difficulty of detecting and differentiating the signal decay caused by different diameters. For the post-mortem case, using the somewhat big *d* = 1 *μ*m and strong Connectom-like acquisition (G = 300 mT/m), we are expecting a signal decay of 0.35%. To be able to statistically identify this signal decay, we would typically need a decay to be at least bigger than ~ 2 standard deviations of the noise (depending on the choice of significance level), which would require SNR ≈ 571.

| **Acquisition parameters** | | **In-vivo** ($D_0 = 2.0 \mu\text{m}^2/\text{ms}$) | | |
| --- | --- | --- | --- | --- |
| $\delta = \Delta$ (ms) | $G$ (mT/m) | $d = 0.5 \mu\text{m}$ | $d = 1.0 \mu\text{m}$ | $d = 2.0 \mu\text{m}$ |
| 10 | 40 | $3.2 \times 10^{-5}$ | $5.2 \times 10^{-4}$ | $8.1 \times 10^{-3}$ |
| 40 | 40 | $1.3 \times 10^{-4}$ | $2.1 \times 10^{-3}$ | $3.3 \times 10^{-2}$ |
| 10 | 300 | $1.8 \times 10^{-3}$ | $2.9 \times 10^{-2}$ | $4.6 \times 10^{-1}$ |
| 40 | 300 | $7.3 \times 10^{-3}$ | $1.2 \times 10^{-1}$ | 1.8 |

Table 1: MR signal **decay** (in percent) for various diffusivities, acquisition parameters, and axon diameters.
| **Acquisition parameters** | | **Post-mortem** ($D_0 = 0.66 \mu\text{m}^2/\text{ms}$) | | |
| --- | --- | --- | --- | --- |
| $\delta = \Delta$ (ms) | $G$ (mT/m) | $d = 0.5 \mu\text{m}$ | $d = 1.0 \mu\text{m}$ | $d = 2.0 \mu\text{m}$ |
| 10 | 40 | $9.8 \times 10^{-5}$ | $1.6 \times 10^{-3}$ | $2.4 \times 10^{-2}$ |
| 40 | 40 | $3.9 \times 10^{-4}$ | $6.3 \times 10^{-3}$ | $9.9 \times 10^{-2}$ |
| 10 | 300 | $5.5 \times 10^{-3}$ | $8.7 \times 10^{-2}$ | 1.3 |
| 40 | 300 | $2.2 \times 10^{-2}$ | $3.5 \times 10^{-1}$ | 5.4 |

Our simulations indicate that dMRI is not very sensitive to the axonal diameter in realistic situations. For example, using optimal in-vivo setting (Connectom strength gradients, very long diffusion pulses, and in-vivo diffusivity) for an axon diameter of 1 micrometer the process only produces a “contrast” of 0.12 % signal decay which is equal to one standard deviation of Gaussian noise with *SNR* ≈ 833. To be able to statistically identify this signal decay, we would typically need a decay to be at least bigger than 2 standard deviations of the noise, depending on the choice of the significance level. To reach such a low noise level would require SNR ≈ 1667. Hence, for realistic SNRs, small diameters cannot be differentiated from the noise level in the image.

### 2.3 Axon diameter limit

To formalize the notion of sensitivity into a workable form using signal decay and SNR, Nilsson et al. [40] introduced the diameter resolution limit (*d*_min_). It is defined as the smallest diameter such that the MR signal decay can be statistically differentiated from no decay (in the limiting case *d* → 0) for a given SNR and choice of significance level for the Z-test (*α*). The decay limit is given by 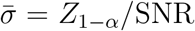. We use eq. 1 to find *d*_min_ corresponding to the decay limit. We use *α* = 0.05 (*Z*_1−0.05_ = 1.645) for the entirety of this work.

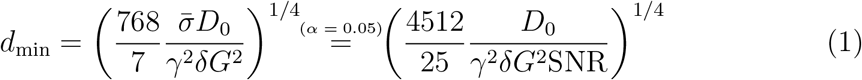

Practically, the main implications of this framework are governed by the exponents of the individual parameters. We can see for instance that halving the diameter limit requires a 4-fold increase in gradient strength or a 16-fold increase in SNR (~ 256 repetitions averaged). Table 2 showcases some values of *d*_min_ for in-vivo and post-mortem diffusivities, a long gradient pulse, various gradient strengths (clinical, Connectom, and small-bore preclinical) for various SNRs. We see that even in the idealized case [40], we obtain *d*_min_ = 2.56 *μ*m for the in-vivo Connectom case at realistic SNR, falling quite short of our minimum target of around 1 *μ*m. At SNR = 164 (~ 5 times higher than baseline, ~ 25 averages), we have 1.77 *μ*m.

**Table 2:** Values of *d*_min_ (*μ*m) (eq. 1) for various parameters at significance level *α* = 0.05 (*i.e.* signal decay stronger than 1.645 standard deviations of the noise distribution). The selected SNRs (164, 65.6, 32.8) correspond to minimum detectable signal decays of 1%, 2.5% and 5%.

| Parameters |  |  | SNR |  |  |
| --- | --- | --- | --- | --- | --- |
| $D_0$ ($\mu\text{m}^2/\text{ms}$) | $\delta = \Delta$ (ms) | $G$ (mT/m) | **164** | **65.6** | **32.8** |
| 2.0 | 40 | 40 | 4.69 | 5.89 | 7.01 |
| 2.0 | 40 | 300 | 1.71 | 2.15 | 2.56 |
| 2.0 | 40 | 1500 | 0.77 | 0.96 | 1.14 |
| 0.66 | 40 | 40 | 3.55 | 4.47 | 5.31 |
| 0.66 | 40 | 300 | 1.30 | 1.63 | 1.94 |
| 0.66 | 40 | 1500 | 0.58 | 0.73 | 0.87 |

In this example, we need tissue with low post-mortem diffusivity and ultra-strong gradients of the strongest preclinical scanner (G = 1500 mT/m) to reach the initial goal of *d*_min_ ≤ 1 *μ*m, showcasing the practical limitations arising from the fourth root scaling in eq. 1.

To visualize the impact of *d*_min_, we plot the spread of recovered diameters in fig. 1. For each diameter between 0.1 *μ*m and 5 *μ*m, we generated 10000 noisy restricted signals and added Gaussian noise with SNR 30 and 300. The signals are generated for realistic in-vivo settings (*D*_0_ = 2 *μ*m^2^/ms) with a Connectom-like acquisition (single “direction/average”, G = 300 mT/m, *δ* = Δ = 40 ms). Both SNRs behave identically with a scale difference and we see that the mean recovered diameter is biased for diameters smaller than *d*_min_. The bias occurs because the average detected diameters become the one corresponding to a signal decay of one standard deviation of the noise. Hence, the result suffers not only from uncertainty but also from systematic bias.

**Figure 1:**
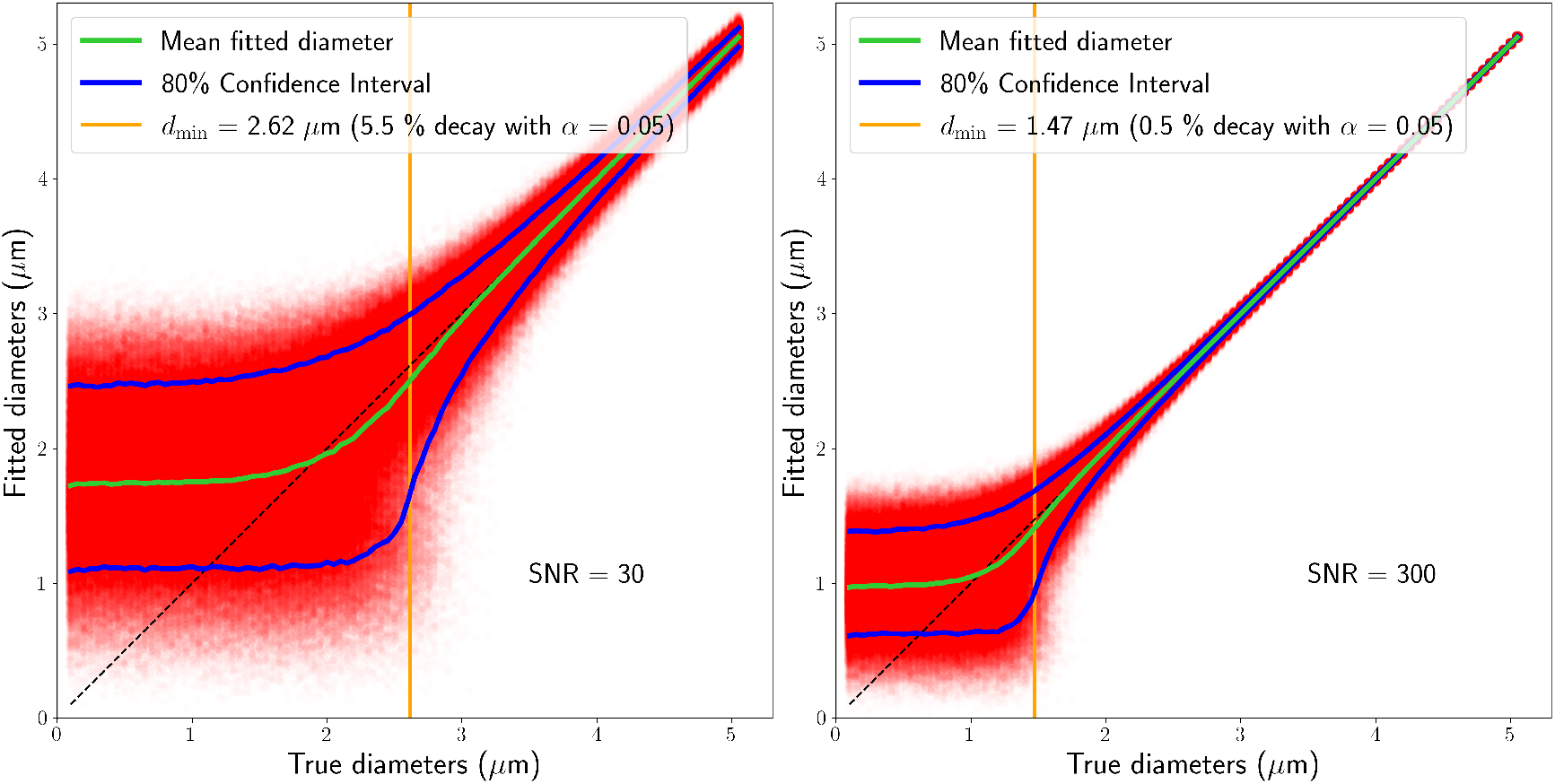
Scatter plot of fitted diameters with mean fitted diameter (green line) and 80% confidence interval (blue lines). For each diameter between 0.1 *μ*m and 5 *μ*m, we generated 10000 noisy restricted signals using eq. 1 and Gaussian noise of SNR 30 (left) and 300 (right). The signals are generated for realistic in-vivo setting (*D*_0_ = 2 *μ*m^2^/ms) with a Connectom-like acquisition (single “direction”, G = 300 mT/m, *δ* = Δ = 40 ms). The orange line corresponds to *d*_min_ using the framework by [40].

It is worthwhile emphasizing what the definition of *d*_min_ truly implies because it is often misunderstood as being the diameter above which fitting will be stable. The formalism of this section is a way to calculate the smallest signal decay *difference* which is statistically differentiable from 0. We can assess if the SNR and acquisition parameters are enough to differentiate two arbitrary diameters, by verifying that their produced signal decay difference is bigger than 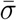. If we set one of those diameters to 0 and we look for the smallest second diameter above the threshold, we get *d*_min_. This minimum diameter only assures us that the distribution formed by repeated noisy signal decay measurements around the true signal decay from a diameter bigger than *d*_min_ doesn’t “overlap significantly” with a signal decay of 0 (*i.e.* less than *α* of the distribution is below 0).

### 2.4 Axon diameter distributions

In previous sections, we focused on the sensitivity of dMRI for axon populations of a single diameter within a voxel. However, the white matter is composed of axons with multiple diameters spanning a large range [1, 37]. Therefore, it is sensible to fit a full distribution of diameters to the measured signal. This strategy can be implemented in multiple ways, such as enforcing a parameterized distribution family such as a gamma distribution over the relative axon counts [7, 11], fitting volume fractions for a binned discretized distribution [16], or by fitting multiple cylinder compartments with diameters as a free parameter. Intuitively, moving from single diameter estimations to any type of distribution will increase the *d*_min_, because adding additional degrees of freedom to a model increases the variance of the fitted parameters [27]. However, the fitting of axonal diameter distributions to dMRI signals is plagued by more than a simple increase to the related *d*_min_, but distribution parameters are unresolvable with PGSE data and there will always be a continuum of equivalent solutions spanning the parameter space.

In this chapter, we show that even the simplest model with multiple diameters has infinitely many completely different solutions for realistic parameters (sec. 2.1). These simulations suggest that any “trend” of different diameters seen in images using such models is not supported by theory and is likely driven by either the regularization terms in the fit or by an effect unrelated to diameter, like noise, errors in the compartment separation or by other types of time-dependent diffusion such as a diffusion signal from the extra-axonal compartment.

When we describe distributions of axon diameters, *P*_axon_(*d*), we refer to distributions over the number of axons (*axon count*) for each diameters inside a voxel. Under the assumption that axons of different diameter have the same proton density, the *spin count* distribution becomes a cylinder volume-weighting of the *axon count* distribution, 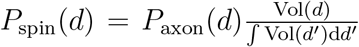. Since the different axons are implicitly assumed to be of the same length inside the voxel, the volume-weighting becomes a cross-section area-weighting 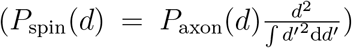. The normalized spin counts are also often referred to as the **volume fractions** of each axon diameter, representing the relative volume of water inside the axons of a given diameter. When the water molecules inside the axons of different diameters have the same magnetic properties (*i.e.* identical T_2_, T_1_, etc), the **signal fractions** are equivalent to the volume-normalized *axon count* distribution. In this study, the conversion between volume and signal fraction only depends on cross-sectional area re-weighting.

In this experiment, we define the simplest distribution, a signal generated from a population of two parallel very big axon diameters in roughly equal proportion (with **signal fractions**: 30% *d*_1_ = 4.5 *μ*m and 70% *d*_2_ = 3.5 *μ*m, equivalent to **volume fractions** of 41.5% and 58.5%) (fig. 2). We then plot the mean absolute difference between this (noiseless) signal and the signals generated for all the other possible configurations.

**Figure 2:**
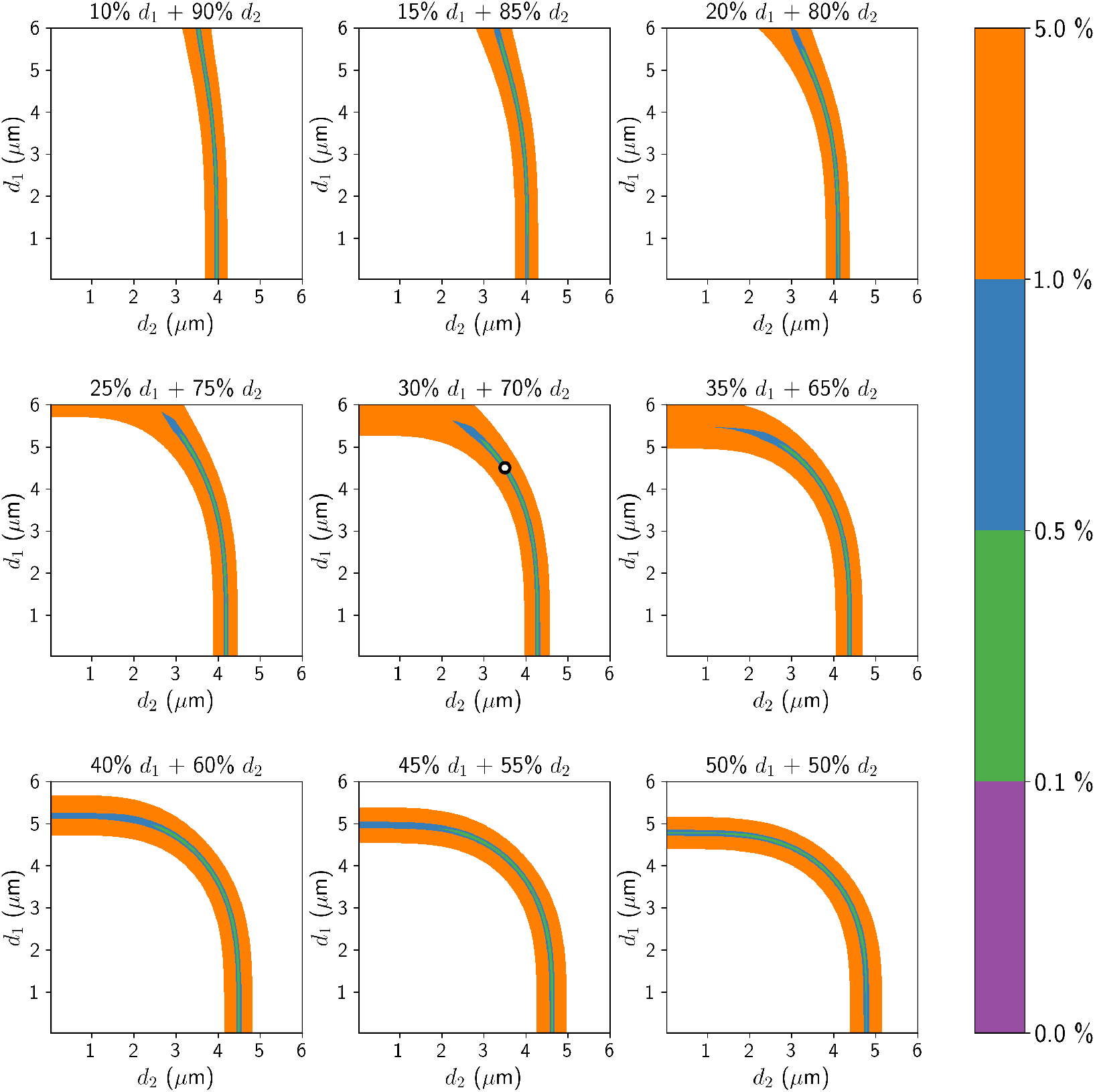
Example of the unresolvability of distribution fitting. The ground truth signal was generated from a combination of 2 parallel cylinders; 30% **signal fraction** with diameter *d*_1_ = 4.5 *μ*m and 70% *d*_2_ = 3.5 *μ*m (shown as white dot in the center plot) with in-vivo diffusivity (*D*_0_ = 2 *μ*m^2^/ms) and a Connectom-like acquisition with three gradient pulse durations (G = 300 mT/m, Δ = 50 ms, *δ* = [30, 40, 50] ms). The parameters were selected so that the smallest diameter was comfortably above the “typical” diameter limit for *δ* = 30 (compared to the limit for SNR = 30, this experiment is noiseless). The 9 subplots represent all combinations of diameters between 0.1 and 6 *μ*m, sliced uniformly at signal fractions between 10% and 50%. The blue “path” corresponds to parameter combinations yielding a signal less than 1% **signal decay** different than the noiseless ground truth. It forms a surface spanning most of the 3D parameter space, rendering any distribution fitting impossible for non-absurd SNR. Section A.4 showcases the same experiment for diameters closer to human axons.

Similarly to how we only used a single “acquisition” (with maximally sensitive Connectom-like parameters) for the single parameter estimation in fig. 1, here we use Connectom-like acquisition parameters with three different gradient pulse durations to mimic the minimal requirements of uniquely fitting a three-parameter model (two diameters and one signal fraction). The acquisition parameters were selected such that they provide sensitivity (long *δ*) and that the biggest individual *d*_min_ is comfortably below the smallest diameter in the ground truth (G = 300 mT/m, Δ = 50 ms, *δ* = [30, 40, 50] ms). This two-cylinder model has a three-dimensional space of possible parameter configurations: the first diameter, the second diameter, and the signal fraction (of the first cylinder). In fig. 2, the parameter space is sliced in the signal fraction direction every 5% and shown as a sequence of 2D plots spanning all pairs of diameters. Regions of solid colors across all slices correspond to regions of the parameter space producing similar signal decay in this noiseless setting. For instance, the blue region corresponds to configurations producing a signal with less than 1% signal decay difference from the ground truth, making them indistinguishable at regular SNR (for example, 1% signal decay corresponds to SNR = 164 for significance level *α* = 0.05). The blue region spans a surface across many unrelated pairs of diameters and signal fractions, showcasing the unresolvability of the simplistic two-diameter distribution under optimal conditions (ground truth perfectly matching the model and no other compartments to disentangle). The axon population diameters were chosen to be very big to highlight the fundamental problem of distribution fitting, for similar figures with smaller diameters, see Sec. A.4 where the effect is amplified.

In fig. 3, we repeat the previous experiment with gamma-distributed axon diameter counts instead of the two-diameter distribution. We generated a signal using a population of cylinders where the *count* for each diameter follows a gamma distribution (shape = 2.25 and scale = 0.4 with peak at 0.5 *μ*m) using the same diffusivities and acquisition parameters as in fig. 2. We show the mean absolute difference between our (noiseless) signal and a signal generated from gamma distributions spanning shapes up to 9 and peak location up to 3 *μ*m. We note that a gamma distribution Γ(*k, θ*) of shape *k* and scale *θ* has its peak at (*k* − 1)*θ* for *k* ≥ 1 (0 otherwise). Regions of solid colors correspond to regions of the parameter space producing a similar signal decay in this noiseless setting. The colored dots in the central parameter space correspond to the signal generated with the corresponding colored distribution (ground truth is red). As was the case with our previous two-cylinder example, we have a wide area of the parameter space generating roughly indistinguishable signals. The four distributions pictured on the sides all produce essentially identical signals for a wide range of distribution shapes.

**Figure 3:**
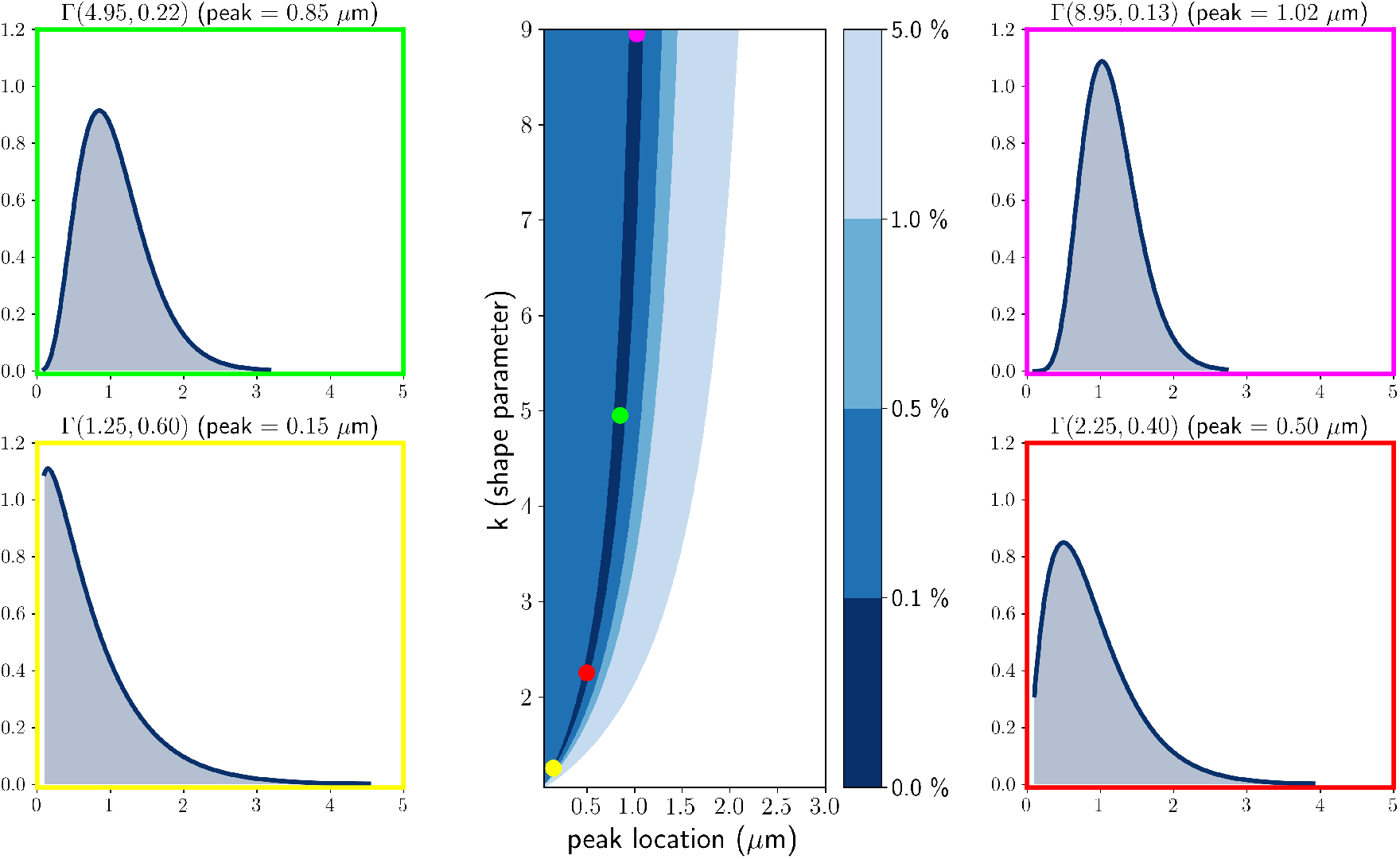
Example of the unresolvability of distribution fitting. The ground truth signal was generated using a gamma distribution of diameter count (shown as a red dot in the center plot) with in-vivo diffusivity (*D*_0_ = 2 *μ*m^2^/ms) and a Connectom-like acquisition with three different gradient pulse durations (G = 300 mT/m, Δ = 50 ms, *δ* = [30, 40, 50] ms). The center plot represents all combinations of shape and peak location characterizing different gamma distributions. The dark blue “path” corresponds to parameter combinations yielding a signal less than 0.1% **signal decay** different than the noiseless ground truth. It forms a path spanning across most of the 2D parameter space, rendering distribution fitting unreliable for non-absurd SNR. The 4 side plots show examples of various gamma distributions from the center plot of wildly different shapes generating roughly indistinguishable signals.

### 2.5 Effective MR diameter

We have shown in the previous section (sec. 2.4) that it seems unfeasible to fit even the simplest distributions. Therefore, we might resort to fitting a single “effective” diameter. When fitting a single parameter over a quantity following a distribution, it is natural that this fitted value will take the form of a central tendency measure of that distribution (a “weighted average”).

In the case of MR axon diameters, two main effects are providing the “weighting”. First, even though we are interested in the distribution of the *axon count*, the signal fractions are weighted by the *spin count*. Under the assumption of uniform intra-axonal proton density, T_2_, same length cylinder for each diameter and no exchange, this manifests itself as a cross-section area weighting, proportional to the 2^nd^ power of the diameter. Secondly, the MR signal is sensitive to the 4^th^ power of the diameter (as seen in eq. 4), adding up an extra heavy tail-weighting effect. Putting it all together, we can define the effective MR axon diameter *d*_eff_ over an arbitrary count distribution of density *P* (*d*) as a function of its moments (eq. 2) [14, 52].

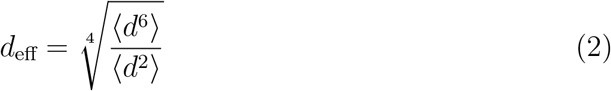

where 〈*d*^*n*^〉 = ∫_*d′*_*P*(*d′*)*d′^n^*d*d′* is the n^th^ moment of the distribution of density *P*(*d*) (See sec A.2 for a simple proof-of-concept derivation). Fig. 4 shows a high match between the effective axon diameter computed from fitting a single diameter over the signal simulated from the distribution (*d*_fit_ in red) and the effective axon diameter derived from direct computation using the moments of the distribution (*d*_eff_ in green) for an example of a human axon diameter distribution from the left and right uncinate/inferior occipitofrontal fascicle taken from [37]. Preliminary post-mortem results [52] indicated a good correspondence between *d*_eff_ estimated from microscopy and dMRI in a rat brain using a complex imaging strategy that properly suppresses non-intra-axonal signals and effects from axon orientations and dispersion.

**Figure 4:**
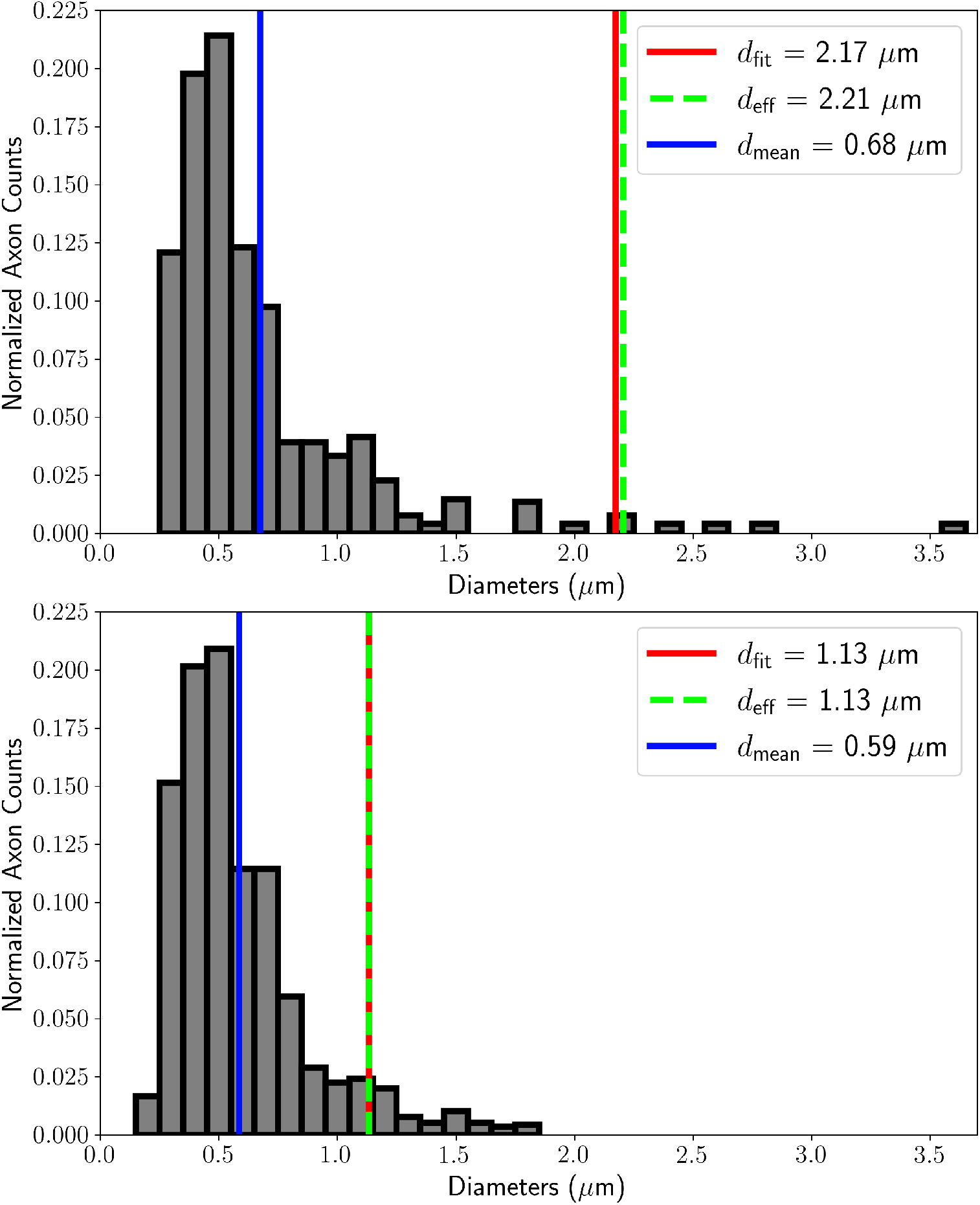
Human axon diameter normalized count distribution taken from Liewald et al.[37] (fig. 9, human brain 1, left and right hemisphere shown as top and bottom respectively). The peak diameter for both distributions is 0.5 *μ*m while the mean diameter *d*_mean_ is around 0.6 *μ*m. The bottom distribution maxes out below 2 *μ*m while the top distribution has a few extra axons in the 2-4 *μ*m range (~ 2.5% of axons by count). This small tail difference heavily affects the effective diameter *d*_eff_ (eq. 2) (doubles it in this case). The fitted MR diameter *d*_fit_ corresponds nicely with *d*_eff_ estimated from the moments of the distribution.

Evidence points toward *d*_eff_ from eq. 2 being an accurate description of the “averaging” process of a typical dMRI sequence over a distribution of axons in the presence of no other signal [47, 52] and a recent in-vivo study [53] assessed the test-retest variability of the metric estimation. However, it is important to keep in mind the limitations of *d*_eff_ as a metric. By the nature of dMRI, *d*_eff_ is extremely weighted toward the tail of the distribution. An Illustration of this phenomenon is shown in figure 4. The two shown axon diameter distributions are fairly similar in terms of mean, peak location (mode) and shape. However, the distribution of the left hemisphere (top plot) comprises an additional ~ 2.5% of large axons, effectively doubling the *d*_eff_ compared to the distribution of the right hemisphere (bottom plot). This difference in the distribution might be due to the small sample size inherent to histology. Such big axon outliers should not be an issue for dMRI where *d*_eff_ is estimated from voxels containing millions of axons. However, when comparing the *d*_eff_ metric between different ROIs or subjects, it becomes impossible to distinguish between situations such as a small global shift toward larger axons or a thicker distribution tail. This is expected when summarizing a complex distribution with only a single metric, instead of the two or three degrees of freedom it requires [46]. Nevertheless, the interpretability of *d*_eff_ is additionally impaired by the heavy tail weighting of its calculation, compared to other single metric distribution summaries, such as the mean. Fig. 5 shows the same axonal diameter distribution taken from Liewald et al. [37] overlapped with densities of multiple families of distributions (gamma, normal, uniform, exponential) with parameters tailored to produce the same theoretical *d*_eff_. The goal is to highlight the large (infinite) number of strikingly different distribution shapes that can produce the same *d*_eff_. The interpretation of *d*_eff_ in its current state will require a very strong hypothesis on the type of distributions or differences that can exist, which is not available in general.

**Figure 5:**
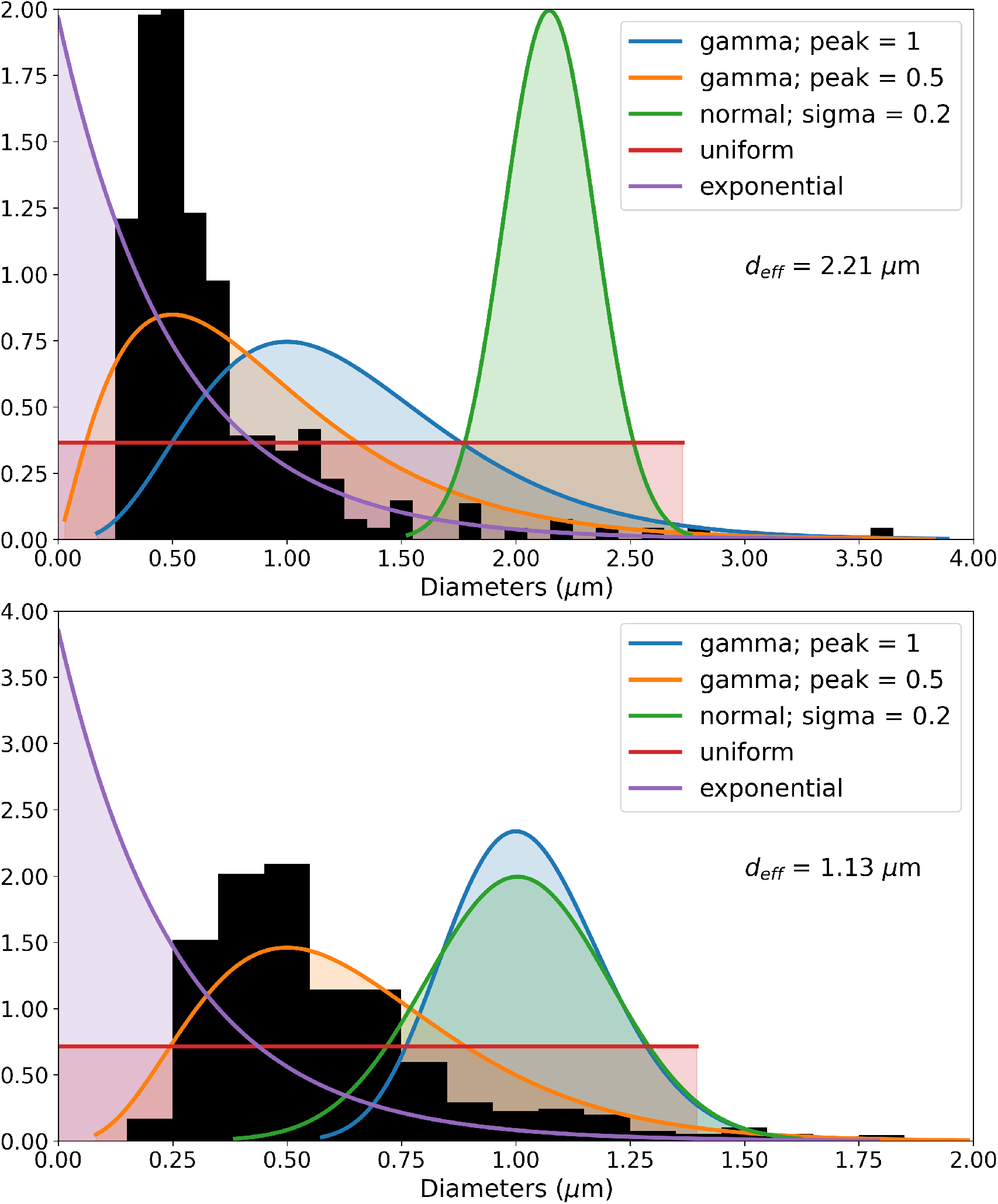
Different families of distributions tuned to produce the same *d*_eff_. The target *d*_eff_ values were computed from the human axons diameter count distribution from Liewald et al.[37] (in black, the discrete counts were converted into a density for visualization). For both hemispheres, we used various families of distribution (restricted to be univariate) to show potential shape variance with identical *d*_eff_.

## 3 Discussion and conclusion

The goal of this work is to showcase the sensitivity limits and the unresolvability of MR axon diameter models from PGSE diffusion-weighted sequences. Those sensitivity limitations affect all dMRI diameter estimation models or techniques, as they are built upon cylindrical compartments and use the same restricted diffusion formulas. In sections 2.2 and 2.3, we have shown how simple computations using realistic in-vivo parameters even with high-end Connectom MR gradient systems generate only very small signal decay with extremely limited sensitivity to relevant axonal diameters. Even the more favorable combination of post-mortem tissue and ultra-strong preclinical gradients does not result in sufficient signal decay to measure realistic axon diameters using diffusion MRI. The problem can be reframed statistically by comparing the signal decay to the noise level with a Z-test and defining a diameter limit. Computing *d*_min_ results in values that are very big compared to relevant axon diameters in the human brain. The effect of this limit was shown with an explicit simulation in fig. 1. In section 2.4, we have shown that fitting a distribution of diameters to the signal results in a multitude of widely different solutions even in the simplest settings. Finally, in section 2.5, we have shown how a distribution of diameters projects itself onto a single fitted effective diameter. The effective diameter can be estimated reliably using advanced hardware and dedicated sequences [52, 53] but still display the same insensitivity to small and medium axons and unresolvability of distribution shape. Nevertheless, the effective diameter is correlated with the biggest axons and differences in these biggest axons might be relevant in some cases such as pathology [53].

We want to emphasize that every result in this work was computed utilizing idealized simulations that were arranged such that any presented limits correspond to a bound on the actual limit on real data. Notably, it has been shown [21, 40] that PGSE is the most sensitive sequence to axon diameters under the assumption of parallel cylinders and perpendicular diffusion gradient. In general, simulations are seen as insufficient to prove or disprove the effectiveness of a method. However, we work on data generated to perfectly follow the general assumptions of the fitted models and we observe that the model fits are still insensitive to the relevant underlying tissue properties. This implies serious doubts that any outputs from those models [7, 3, 25, 16, 24] applied to real data represent the underlying axonal diameter information, no matter how visually appealing they might seem. We suspect that previous, apparently plausible results arise from some combination of effects outside of the model assumptions that are projecting themselves onto the model parameters in a complex way. Admittedly, some parameters can be reliably fitted to the data, such as the *effective diameter* [52, 53], if an appropriate sequence and method are used to suppress the extra-axonal signal contribution. As we have demonstrated, such indices are insensitive to small and medium axons, and to the shape of the axon diameter distribution but they might still provide useful information to study cases principally influenced by big axons.

Hence, any claim of the infeasibility of axon diameter measurement based on the employed simulations automatically translates to the infeasibility of axon diameter measurements based on real data acquired with similar parameters. Our simulated data were generated (I) purely from intra-axonal signals and (II) perpendicular to the main orientation. In a multi-compartment model where the extra-axonal signal has to be fitted, (III) there will be residual fitting errors from the extra-axonal compartment contaminating the already tiny intra-axonal signal decay, increasing the effective *d*_min_. For example, a typical extra-axonal tensor compartment in the WM with a perpendicular diffusivity of 0.3 *μ*m^2^/ms produces a signal decay of ≈ 72.3% for acquisitions parameters *δ* = Δ = 10 ms and *G* = 300 mT/m. If only 1% of this signal decay (i.e. 0.723% total signal decay) is instead erroneously considered as restricted compartment decay fitted with *D*_0_ = 2 *μ*m^2^/ms, it would be equivalent to a cylinder with a diameter 2.25 *μ*m

In the simulations, we considered that the typical white matter SNR from an MR acquisition using Connectom gradients was driven only by the intra-axonal compartment. (IV) However, in reality, the intra-axonal volume fraction comprises less than 50% of the total volume in dense parallel fiber regions such as the corpus callosum and less in deep white matter [15]. This discrepancy (at least) halves the measured intra-axonal signal decay, thereby additionally increasing the effective *d*_min_. (V) Moreover, uncertainties in the estimation of the fiber orientation will additionally bias the apparent diameter because the restricted diffusion model will be fitted to the elongated elliptical cross-section. (VI) Unaccounted orientation dispersion for multi-compartment models will make estimation essentially impossible as shown in Nilsson et al. [40]. Considering all those sources of bias, it is clear that the already small signal decay caused by the restricted diffusion inside axons is essentially unattainable with such multi-compartment models.

An important message from eq. 1 and tables 1–2 are the scaling powers of the parameters. They are such that the sensitivity problem cannot be fixed using more powerful gradient systems. Even extreme cases such as going from in-vivo Connectom-like (G = 300 mT/m) acquisitions at normal SNR, to post-mortem measurements with ultra-strong preclinical gradients (G = 1500 mT/m) and 5 times better SNR (25 averages) only decreases the *d*_min_ from 2.56 to 0.58 *μ*m (around 4.4 times better). This new value is barely enough to be sensitive to the peak of the diameter distribution in the best case. If we consider all the idealized assumptions from the diameter limit formula, it is likely not sufficient.

There are many misconceptions in the literature about the difficulty of going from single diameter fitting to multiple diameters or a distribution. The “intuition” that errors in the fitted distribution will be distributed around the true solution fails spectacularly, even in the absolute simplest case of a signal from two axonal compartments with big diameters and no source of possible confounds as seen in fig. 2 and section A.4. A commonly seen argument is to limit the distribution fit at some *d*_min_ best-case value and claim that the resulting distribution must be valid because we are sensitive to these bigger diameters. Ignoring the value of *d*_min_, we can focus on what it fundamentally attempts to do, putting a limit on the minimal signal decay that can be statistically seen above the noise. To highlight this previous point, fig. 2 shows where configurations such as (35% 4.95 *μ*m + 65% 2.9 *μ*m), (30% 4.5 *μ*m + 70% 3.5 *μ*m), (100% 3.85 *μ*m) and (45% 0.1 *μ*m + 55% 4.6 *μ*m) produced signal with [0.1, 0.5]% signal decay difference. Such a small decay requires SNR ∈ [329, 1645] for detection at optimal in-vivo Connectom-like settings, which correspond to a *d*_min_ ∈ [0.96, 1.44] *μ*m, showing the disconnection between the limits of distribution fitting and direct *d*_min_ computation.

With the complexity of real axonal diameter distributions and the apparent impossibility of reliably fitting a distribution, working with the effective diameter *d*_eff_ seems to be the most promising avenue. When using an advanced acquisitions strategy to negate the non-intra-axonal signal [52], we can accurately estimate *d*_eff_. However, *d*_eff_ is not a well “behaved” metric for comparisons involving different shapes of axonal diameter distributions, such as between subjects or different brain areas. For a complete analysis, we would potentially need to develop a new non-Stejskal-Tanner diffusion sequence producing a slightly different weighting of the distribution to allow some shape disentangling. In its current state, the metric *d*_eff_ cannot differentiate fundamentally different situations such as a small diameter increase of all axons versus a large diameter increase from a small proportion of the axon population. In pathological cases where the very few extra-large axons (or possibly glial cells) are affected, the effective diameter might provide additional information [53]. However, for many other purposes that require the full distribution, in particular for neuronal modeling, it is unsuitable.

The conduction velocity of myelinated axons is strongly impacted by axon diameter [54, 20]. The results presented in this work seem to indicate that direct conduction velocity estimations from dMRI are unfeasible since they rely entirely on axon diameter results suffering from sensitivity limitations. The missing sensitivity of small to medium axon diameters translates into the overestimation of conduction velocities [28, 32]. Similarly, any error in compartment separation or main orientation estimation is likely to a cause large bias in the diameter estimation. Under the most simplistic model, the conduction velocity of myelinated axons is a linear function of the outer axon diameter (*i.e.* including myelin sheath) [31], as well as the inner axon diameter under equal g-ratio. Under this relation, the conduction velocity estimated using *d*_eff_ is heavily weighted toward the conduction velocity of the largest axons, following the same pattern as Eq. 2 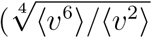 for conduction velocities *v*) and suffering from all the same limitations.

An apparent oversimplification throughout this work concerns how SNR and the number of samples are chosen. For example, in fig. 1, our 1D approach is equivalent to generating the signal for a single gradient direction perpendicular to the cylinder. Similarly, we chose three *δ* for fig. 2 *i.e.* equal to the number of free parameters for the two-cylinder model. If one has a real sample containing only identical parallel cylinders, the knowledge of the orientation would not be present and hundreds of directions would be sampled across multiple values of *δ* and *G*. It is hard to define a single value representing the SNR gain going from one data point with perfect alignment and with maximal sensitivity to hundreds of data points with varying sensitivity, extra parameters to fit, etc. If we take instead 100 repetitions of the optimal measurement and ignore the unknown orientations, we get an upper bound of 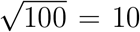 times better SNR which corresponds to a 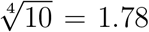 times smaller *d*_min_. A more realistic upper bound is to include the estimation of the direction as two extra free parameters and frame the data as 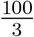 repetitions of three optimal measurements; 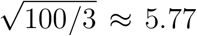 times better SNR which corresponds to a 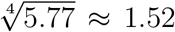 times smaller *d*_min_. This view becomes increasingly complex as we add more parameters and start taking into account how different measurements have non-equal sensitivity to each of the estimated parameters. Since there is a 8^th^ root scaling of *d*_min_ versus additional averaging (functional form of diameter versus signal decay is 4^th^ power and SNR versus averages is 2^nd^ power in the best case), we feel that results on a minimal number of data points are sufficiently relevant.

An interesting topic we did not mention so far is the time-dependence of the extra-axonal space diffusion [14, 18, 26, 35, 42]. Previous attempts to model axonal diameters assumed that all the time-dependent diffusivity portions of the signal were due to intra-axonal restricted diffusion. Recent work has highlighted a mechanism by which the extra-axonal space can also produce signals with time-dependent diffusivity. Indeed, the spacing of the restricting barrier in the extra-axonal compartment tends to be larger than typical axon diameters at relevant time-scales. This has the effect of producing a larger signal decay than the intra-axonal restricted compartment for a given acquisition scheme and producing a time-dependent diffusivity when varying Δ. We briefly show in section A.3 how this extra-axonal time-dependence could contribute to the axon diameter overestimation seen in literature [3, 5, 23, 29, 30].

In summary, our results show that the MR-based assessment of axonal diameters is methodologically infeasible. Our simulations under ideal conditions demonstrate that diffusion-weighted MRI with current and foreseeable future hardware is not capable of performing axonal diameter measurements in biologically relevant dimensions. The inability to measure axonal diameters is not a matter of the biophysical model choice but rather stems from the missing contrast of the intra-axonal tissue fraction. Under realistic, less ideal measurement conditions, the feasibility of such measurements is even further reduced. We show that frequently shown “known” variations of axonal diameter across structures such as the corpus callosum might also be explained with time-dependent diffusion of the extra-axonal tissue fraction. Therefore, previous measurements and model fitting results rather represent a characterization of the extra-axonal space than a measure or representation of the axonal diameter. Our manuscript further investigates recent descriptions of axonal diameters using a projection on an “effective diameter”. Our simulations show this representation can be strongly affected by small changes in the distribution tail and does not allow to draw any unambiguous conclusions about the actual distribution of diameters.

Given the immense methodological difficulties of MR axonal diameter measurements, we suggest including the time dependence of extra-axonal diffusion in the quantitative description of the microstructure of white matter in future studies (as in [18]). In connection with an independent measure of tissue myelination, this time dependency may provide an indirect approach to estimate the outer axonal diameter. Multidimensional dMRI measurements [50] may help to describe the extra-axonal space due to a reduced degeneracy of associated microstructural models. This may open a doorway to a quantitative study of brain microstructure using diffusion MRI.

## Acknowledgement

MP is supported by a scholarship (PDF-502732-2017) from the Natural Sciences and Engineering Research Council of Canada (NSERC). MP and CE are supported by the Priority Program 2041 (SPP 2041) “Computational Connectomics” of the German Research Foundation (DFG).

## A Appendices

### A.1 Insensitivity of Δ to axon diameter

There is some misunderstanding in the literature concerning the impact of varying Δ to probe axon diameter. Intuitively, the dMRI signal is created by the dephasing of spins due to their displacement. For Δ to play a role in the measured restricted signal, we need to be in a short enough *δ* time regime. In the long time regime (wide pulse regime), by the end of the gradient application, most spins have interacted strongly with the axonal wall and their positions are mostly de-correlated from their initial position; the maximal signal decay has been reached and changing the gradient spacing Δ will not change anything. In the range of relevant parameter values (see sec. 2.1), it is simple to numerically show this phenomenon. Fig. 7 shows the signal decay computed from eq. 3 for all physically plausible (Δ, *δ*) pairs in Δ ∈ [10, 50] ms and *δ* ∈ [10, 50] ms for various axon diameters for in-vivo Connectom-like settings. The respective signal decay depends strongly on the diameters, however, there is no perceptible difference for different Δ at the same *δ*. The same results can be achieved by Monte-Carlo spin diffusion simulation (see Fig. 6).

**Figure 6:**
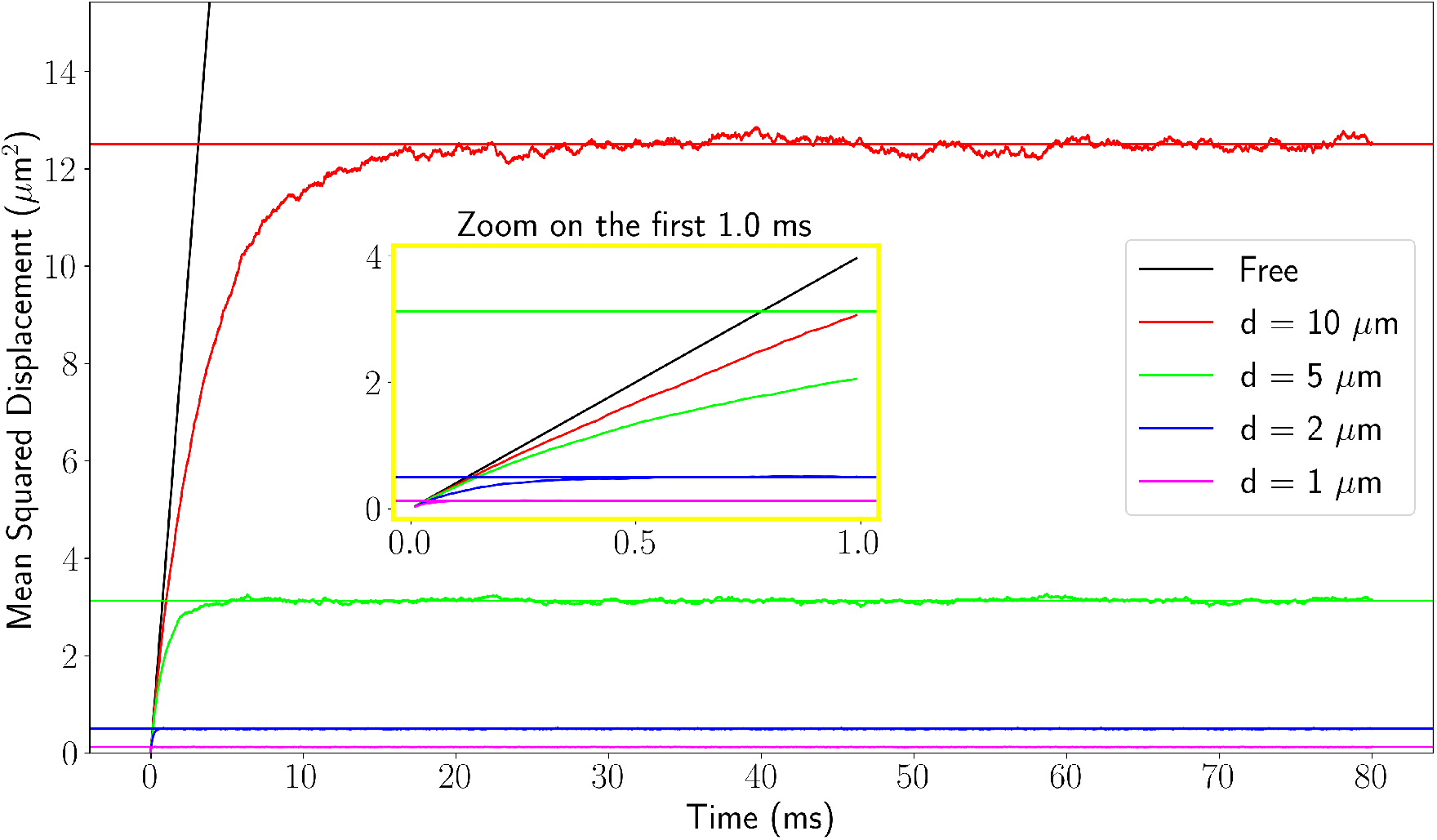
Mean squared displacement (MSD) for one direction from 2D Monte-Carlo simulation for free diffusion and restricted diffusion inside circles of different radii using *D*_0_ = 2 *μ*m^2^/ms. The horizontal lines show the long time limit MSD for each diameter. The center plot is a zoom on the first millisecond where we see that even the relatively large 2 *μ*m diameter circle reaches long time regime quicker than any sufficiently strong diffusion gradient can be applied (*δ*_min_ ≥ 5 ms).

**Figure 7:**
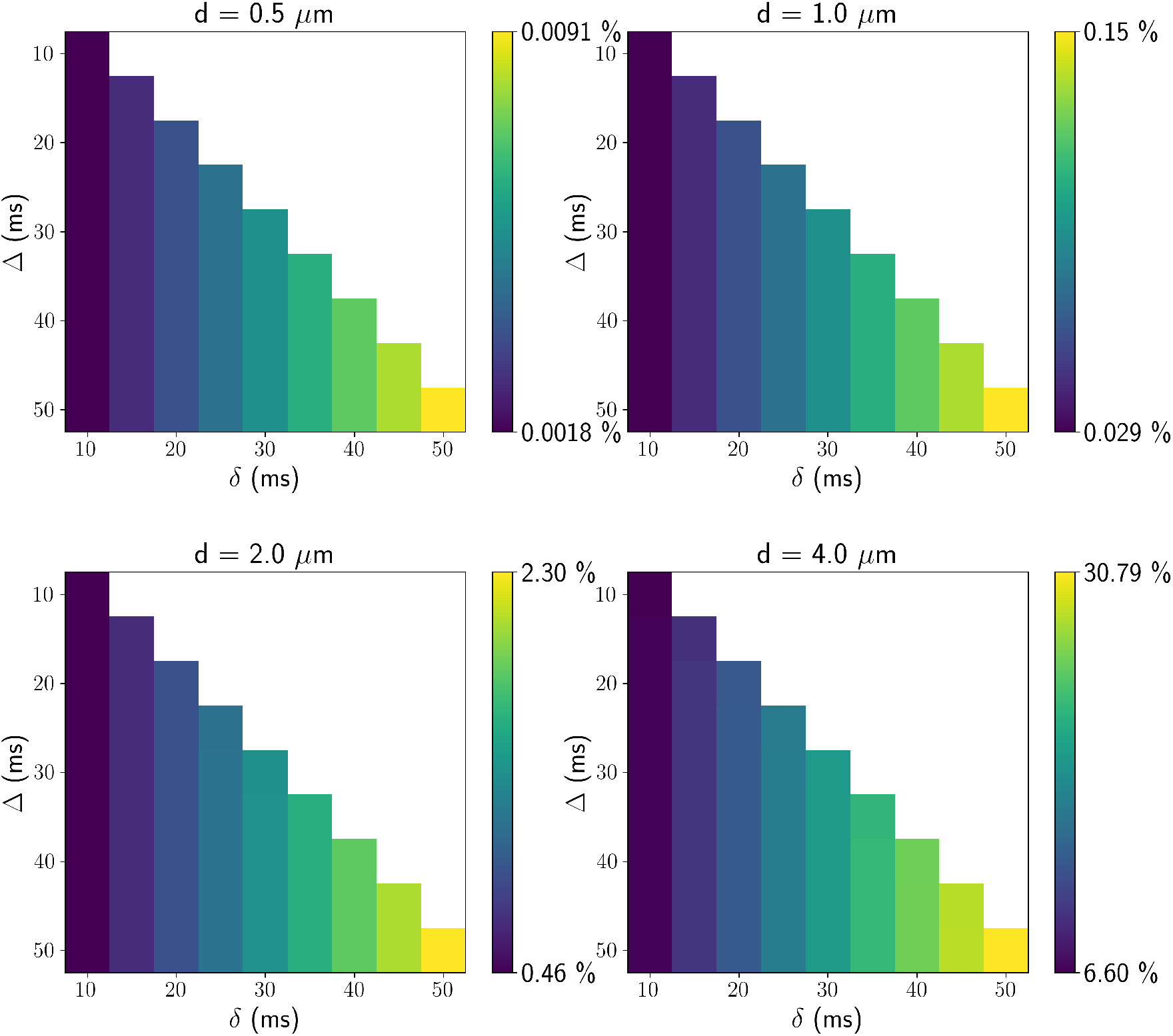
Noiseless MR signals from eq 3 for various (Δ, *δ*) and axon diameter (*d*). The signals were simulated for *G* = 300 mT/m and *D*_0_ = 2.0 *μ*m^2^/ms. We note that different Δ (y-axis) doesn’t modify the signal in any appreciable way.

Another way to demonstrate this result is to derive the rough form of the signal equation from spin dephasing [35]. We have applied gradient *g* and pulse width *δ*. In the long time regime (wide pulse regime), we have *δ* ≫ *t*_*c*_, *t*_*c*_ being the characteristic correlation time of the cylinder (*t*_*c*_ ~ *d*^2^*/D*_0_). We will first calculate the phase *ϕ*_1_ accumulated by spins within a time window of *t*_*c*_ (where the Gaussian phase approximation applies [39]) and then compute the total phase *ϕ* accumulated as a sum of *N* ~ *δ/t*_*c*_ uncorrelated contributions. Within one short step, phase is accumulated linearly proportional to the applied gradient and spin displacement, *ϕ*_1_ ~ *gdt*_*c*_. We now compute the signal using 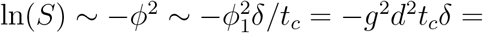 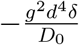. The recovered equation form corresponds to the Neuman long-time limit up to a constant and is independent of Δ and the initial position (it implicitly vanished by considering a displacement of *d* for a time-step of *t*_*c*_ in *ϕ*_1_).

The Van Gelderen formula (Eq. 3) for restricted diffusion is derived using the Gaussian phase approximation (GPA), which assumes a Gaussian distribution for spin dephasing. The GPA is guaranteed to hold for very short or very long diffusion time [39] and is empirically accurate for simple shapes of boundaries such as circles. It is a fast converging series, typically reaching precision errors smaller than 10^−7^ with only 10 terms. Often, instead of truncating Eq. 3, we use the Neuman long-time limit formula (Eq. 4) for simplicity. It is typically numerically indistinguishable from Eq. 3 for the relevant parameter range. The Neuman formula is defined assuming Δ = *δ*, but it can nonetheless be used in the case of the relevant parameter range where the restricted diffusion signal is insensitive to Δ

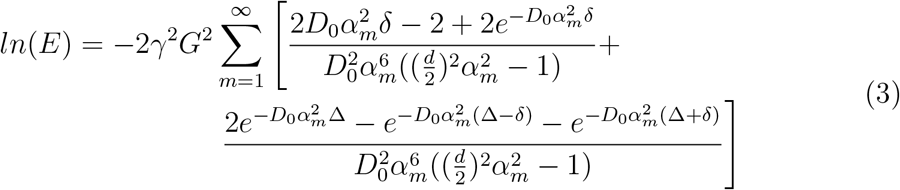

where E is the normalized diffusion signal, *γ* is the proton gyromagnetic ratio, *G* is the diffusion gradient amplitude, *D*_0_ is the unrestricted diffusivity in the cylinder, Δ is the diffusion gradient separation, *δ* is the diffusion gradient duration, *d* is the diameter of the cylinder, *α*_*m*_ is the m^th^ root of the equation 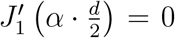, where *J′*_1_ (·) is the derivative of the Bessel function of the first kind.

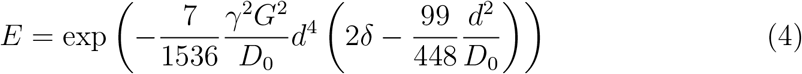

### A.2 Effective diameter derivation

We give a simple derivation of the effective diameter (similar derivation can be found in [14, 52]). The normalized MR signal as a function of *d* with all other parameters fixed is

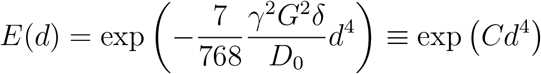

for some fixed constant *C*. We compute the volume fraction normalized signal *E*_*P*_ for diameter **counts** following a distribution of density *P*(*d*). We use the approximation *E*(*d*) ≈ 1 + *Cd*^4^ from the truncated Taylor series of *exp*(·).

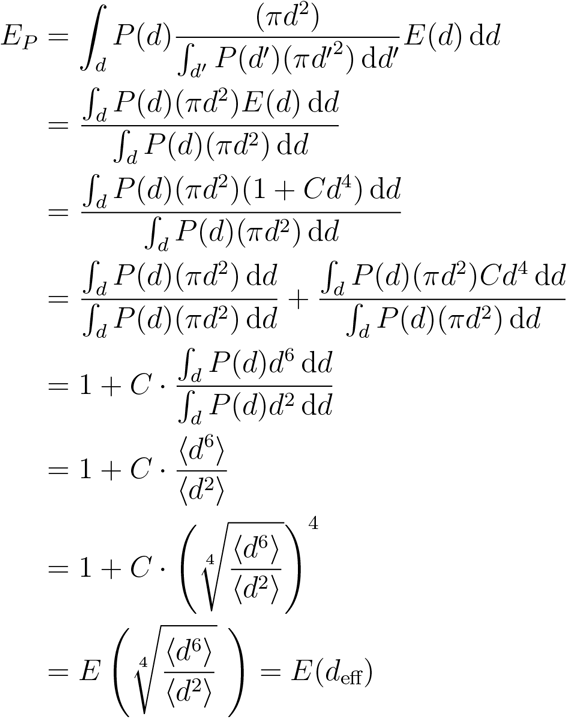

### A.3 Extra-axonal time-dependent diffusivity

It has been shown that the extra-axonal compartment can exhibit time-dependent diffusivity [14, 26, 35, 42]. It arises from the disorder created by the irregular packing of axons of varying diameters. The “disorder strength” is characterized by the parameter *A* and has been empirically estimated in [14] to be 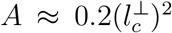 where 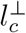 is the fiber packing correlation length at which diffusion is restricted in extra-axonal space. Two models of perpendicular diffusivity as a function of (Δ, *δ*) are described in [35]; 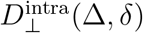 assuming that all the time dependence in the diffusivity arises from intra-axonal space, 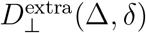 assuming that all the time dependence in the diffusivity arises from the extra-axonal space.

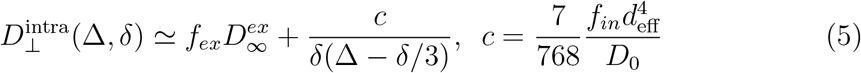

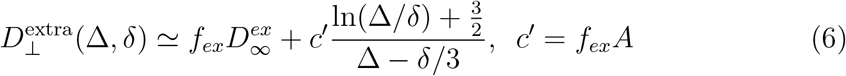

with extra-axonal volume fraction *f*_*ex*_, intra-axonal volume fraction *f*_*in*_ = 1−*f*_*ex*_, long time (Δ → ∞) extra-axonal diffusivity 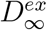, bulk diffusivity *D*_0_, and disorder strength parameter *A*.

Evidence on a few subjects suggests that the extra-axonal time-dependence dominates the intra-axonal time-dependence [18, 35]. This was shown by fitting both eq. 5 and 6 to data acquired with fixed *δ* = 20 ms and multiple Δ ∈ [26, 100] ms to comparable goodness-of-fit. The fitted parameters were then used to predict the signal values of a second acquisition using Δ = 75 ms and multiple *δ* ∈ [4, 45] ms, where the extra-axonal model obtained good predictions and the intra-axonal model failed. Since most axon diameter estimation methods assume static values for the extra-axonal diffusivity, if the time-dependence in the signal is dominated by extra-axonal effects, the estimated diameters will be large and mostly unrelated to the effective diameter *d*_eff_. To showcase this effect, we equated eq. 5 and 6 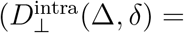 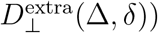 and isolated *d*_eff_. We used the typical value of *D*_0_ = 2 *μ*m^2^/ms and fixed 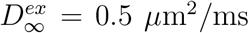 (fitted values in [35] inside [0.38, 0.6] *μ*m^2^/ms). We use *f*_*ex*_ ∈ [0.25, 0.75] and *A* ∈ [0.25, 2], giving us *f*_*ex*_*A* ∈ [0.0625, 1.5] compared to the reported values in [35] inside [0.24, 0.56]. We generated the *“fake” d*_eff_ for all physically plausible combinations of Δ ∈ [5, 100] ms and *δ* ∈ [5, 50] ms. We observe effective diameter between 2 *μ*m and 9.5 *μ*m, with most diameters above 6 *μ*m in the configurations (*f*_*ex*_ = 0.5 and *A* = [0.5, 1]) closest to results from [35].

The well-known “small-big-small diameter pattern” observed in the corpus callosum with histology and “reproduced” with big overestimation by axon diameter estimation methods ([3, 16, 23, 38]) can potentially be explained by this presented effect [18]. A brain area with a higher mean diameter is likely to also have an increased 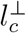 for random circle packing; if the diameter distribution is uniformly shifted up, the packing keeps the same relative efficiency and the individual inter space grows, alternatively, if a few more big axons are present, it increases the diameter heterogeneity and the packing efficiency tend to go down, creating more extra-axonal space. In any case, *f*_*ex*_*A* increases, and the *“fake” d*_eff_ follows in the setting of fig. 8. However, the extra-axonal model parameters still contain some information about the *outer* diameter distribution, but it is complexly tangled with axon packing.

**Figure 8:**
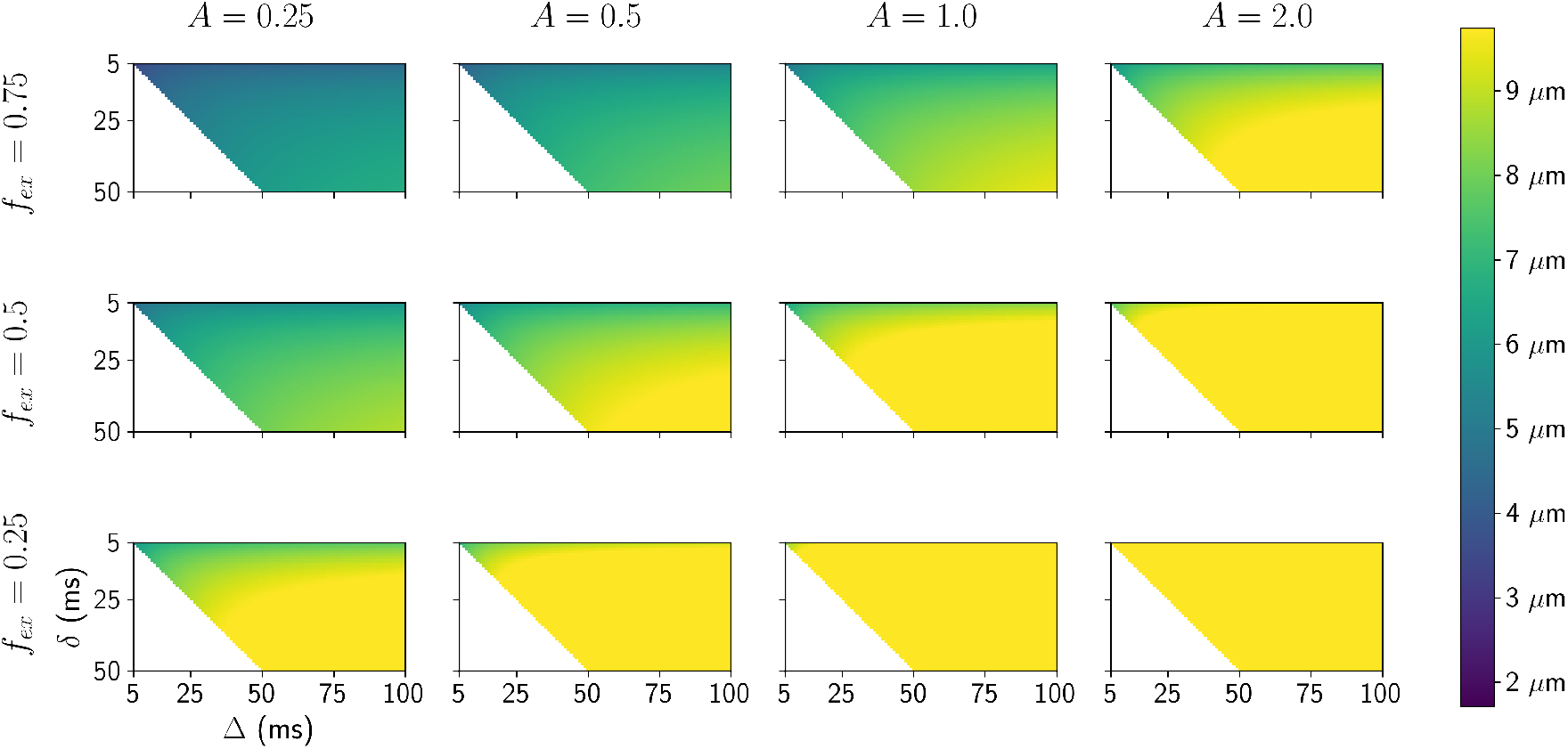
Signal generated using the extra-axonal time-dependence formula eq 6 and effective diameters fitted using eq 5.

**Figure 9:**
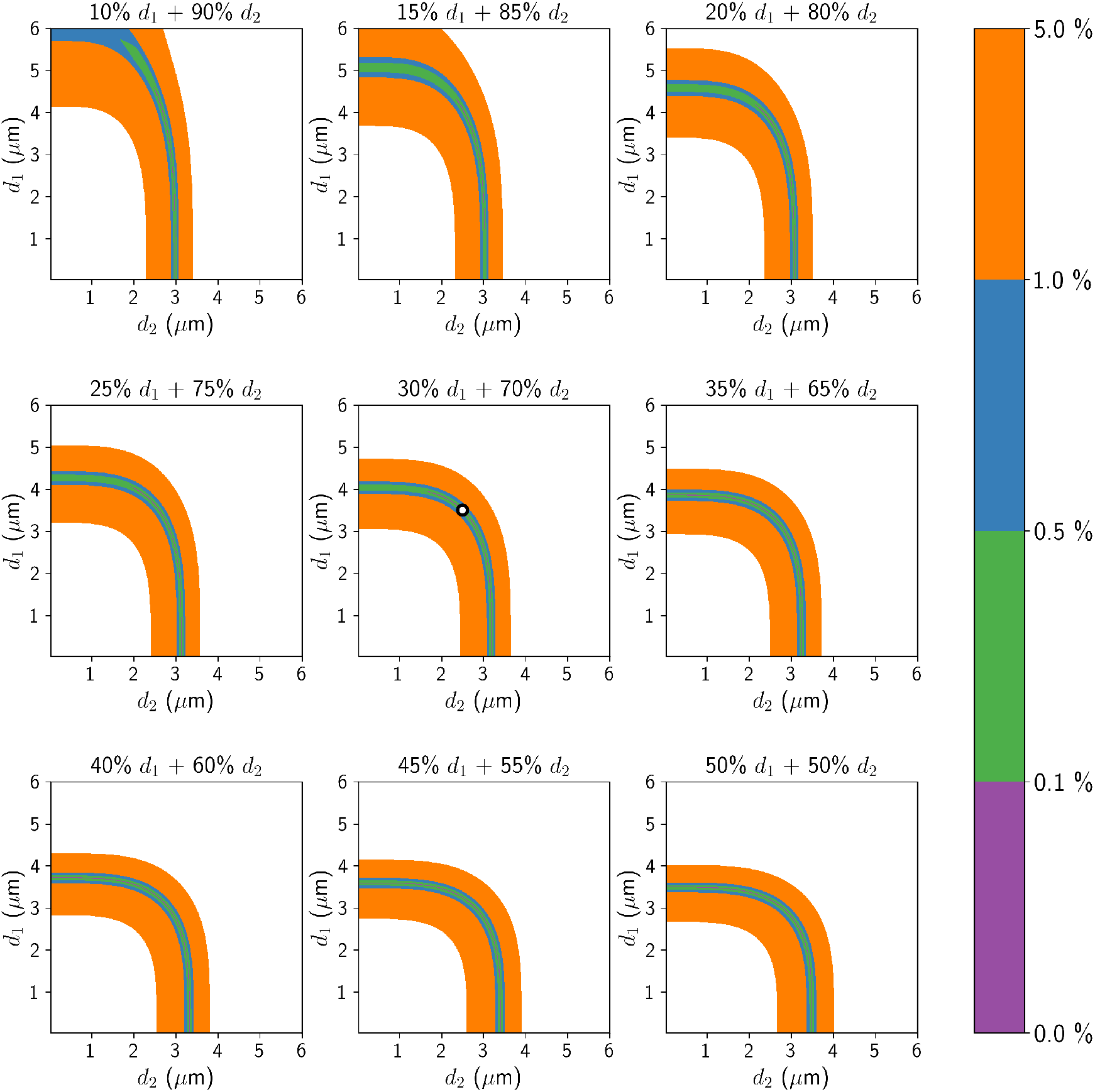
Example of the unresolvability of distribution fitting. The ground truth signal was generated from a combination of 2 parallel cylinders; 30% **signal fraction** with diameter *d*_1_ = 3.5 *μ*m and 70% *d*_2_ = 2.5 *μ*m (shown as white dot in the center plot) with in-vivo diffusivity (*D*_0_ = 2 *μ*m^2^/ms) and a Connectom-like acquisition with three different gradient pulse durations (G = 300 mT/m, Δ = 50 ms, *δ* = [30, 40, 50] ms). The parameters were selected so that the smallest diameter was comfortably above the “typical” diameter limit for *δ* = 30 (compared to the limit for SNR = 30, this experiment is noiseless). The 9 subplots represent all combinations of diameters between 0.1 and 6 *μ*m, sliced uniformly at signal fractions between 10% and 50%. The blue “path” corresponds to parameter combinations yielding a signal less than 1% **signal decay** different than the noiseless ground truth. It forms a surface spanning most of the 3D parameter space, rendering any distribution fitting impossible for non-absurd SNR.

**Figure 10:**
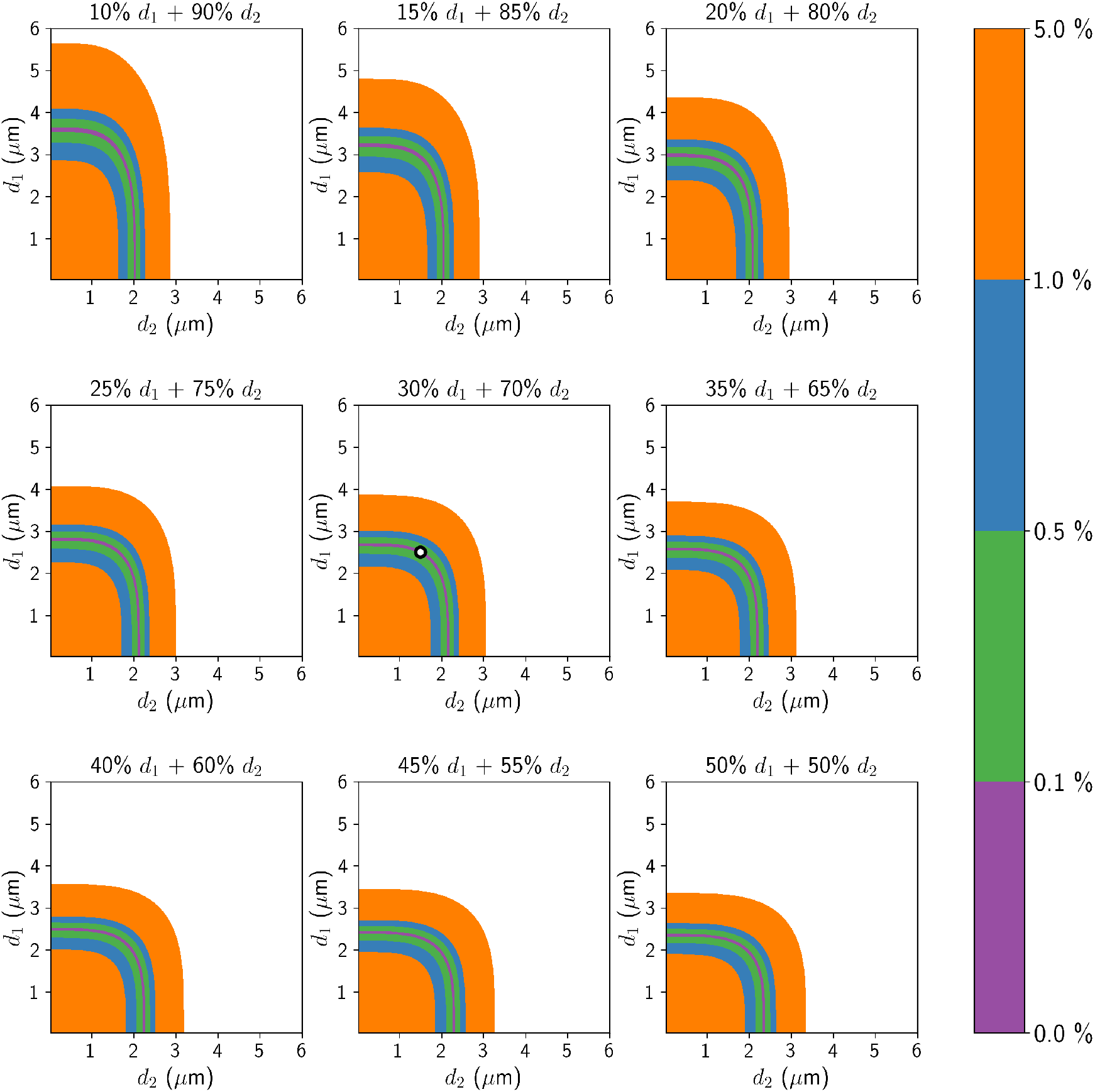
Example of the unresolvability of distribution fitting. The ground truth signal was generated from a combination of 2 parallel cylinders; 30% **signal fraction** with diameter *d*_1_ = 2.5 *μ*m and 70% *d*_2_ = 1.5 *μ*m (shown as white dot in the center plot) with in-vivo diffusivity (*D*_0_ = 2 *μ*m^2^/ms) and a Connectom-like acquisition with three different gradient pulse durations (G = 300 mT/m, Δ = 50 ms, *δ* = [30, 40, 50] ms). The parameters were selected so that the smallest diameter was comfortably above the “typical” diameter limit for *δ* = 30 (compared to the limit for SNR = 30, this experiment is noiseless). The 9 subplots represent all combinations of diameters between 0.1 and 6 *μ*m, sliced uniformly at signal fractions between 10% and 50%. The blue “path” corresponds to parameter combinations yielding a signal less than 1% **signal decay** different than the noiseless ground truth. It forms a surface spanning most of the 3D parameter space, rendering any distribution fitting impossible for non-absurd SNR.

### A.4 Two-diameter distributions

We show more examples of fitting a two-diameter model with smaller, more realistic diameters. In fig. 2, we used a combination of enormous diameters (**signal fraction**, 30% *d*_1_ = 4.5 *μ*m and 70% *d*_2_ = 3.5 *μ*m) to highlight the effect of having a distribution over the lack of sensitivity of the realistic state-of-the-art acquisition scheme. We now show results for (30% *d*_1_ = 3.5 *μ*m and 70% *d*_2_ = 2.5 *μ*m) and (30% *d*_1_ = 2.5 *μ*m and 70% *d*_2_ = 1.5 *μ*m), where the ambiguity over the diameters is amplified for the same sampling scheme.

